# Metabolic signature in nucleus accumbens for anti-depressant-like effects of acetyl-L-carnitine: An *in vivo* ^1^H-magnetic resonance spectroscopy study at 14 T

**DOI:** 10.1101/690768

**Authors:** Antoine Cherix, Thomas Larrieu, Jocelyn Grosse, João Rodrigues, Bruce McEwen, Carla Nasca, Rolf Gruetter, Carmen Sandi

## Abstract

**Background:** Emerging evidence suggests that hierarchical status may provide vulnerability to develop stress-induced depression. Energy metabolism in the nucleus accumbens (NAc) was recently related to hierarchical status and vulnerability to develop depression-like behavior. Acetyl-L-carnitine (LAC), a mitochondria-boosting supplement, has shown promising antidepressant-like effects opening promising therapeutic strategies for restoring energy balance in depressed patients. Here, we investigated the metabolic impact in the NAc of antidepressant LAC treatment in chronically stressed mice.

**Method:** Mice were characterized for emotional behaviors and social rank. They were then exposed to chronic restraint stress (CRS) for 21 days and subsequently tested in a social behavior (SB) test. A group of mice was also given LAC supplementation during the 7 last CRS days. Mice were then tested in the SB and forced swim tests (FST) and scanned *in vivo* using ^1^H-magnetic resonance spectroscopy (^1^H-MRS) to quantitatively assess the NAc neurochemical profile.

**Results:** Dominant, but not subordinate, mice showed behavioral vulnerability to CRS. In the NAc, dominant mice showed reduced levels of several energy-related metabolites. LAC treatment counteracted stress-induced behavioral changes in dominant mice, and normalized levels of taurine, phosphocreatine, glutamine and phosphocholine in the NAc. No major accumbal metabolic changes were observed in subordinate mice.

**Conclusion:** High social rank is confirmed as a vulnerability factor to develop chronic stress-induced depressive-like behaviors. We reveal a metabolic signature in the NAc for the antidepressant-like effects of LAC in vulnerable mice, characterized by restoration of stress-induced alterations in neuroenergetics and lipid function.

## Introduction

Depression is among the leading causes of disability worldwide, which reflects the current lack of understanding of its underlying mechanisms (1, 2). Metabolic alterations are emerging as key etiological factors for the development of neuropsychiatric disorders, including depression (3–5). The strong reliance of the brain on high energy consumption would make it particularly vulnerable to metabolic alterations (3). In addition, chronic stress has a strong capacity to trigger and exacerbate depression (6, 7) and impinges metabolic-costly neuronal adaptations in structure and function (6, 8, 9). Accordingly, stress-associated depletion of brain’s energy resources could lead to impaired neuronal plasticity underlying depression (10, 11).

However, not all individuals are equally affected by stress (9, 12, 13); while some individuals show a high vulnerability to develop depression, others endure resilience following stress exposure (13, 14). It remains unclear which factors provide resilience to stress in certain individuals and what are the underlying mechanisms (15, 16). In addition to the great predictive power of high anxiety trait in defining stress vulnerability (13, 14, 17, 18, 20; for reviews, see Sandi and Richter-Levin, 2009; Russo et al., 2012 and Weger and Sandi, 2018), epidemiological, clinical and animal work point to a link between social hierarchies and depression (15).

Recently, in the C57BL/6J inbred mouse strain, we found that dominant animals were more susceptible to display social avoidance following exposure to chronic social defeat stress (CSDS), while subordinate mice were not affected (8). Data from ^1^H-magnetic resonance spectroscopy [^1^H-MRS; one of the few non-invasive methods that can provide direct information on brain metabolism *in vivo* (20)] revealed a relationship between the metabolic profile of the nucleus accumbens [NAc; a hub brain region for the regulation of motivated behaviors (21) implicated in the pathophysiology of depression (22)], social rank and vulnerability to stress. Thus, while under basal conditions subordinates showed lower levels of energy-related metabolites than dominants, it was only the subordinate/resilient group that displayed increased metabolite levels following CSDS (8). These observations suggested that metabolic targeting may be an optimal treatment intervention.

Acetyl-L-carnitine (LAC) has been recently shown to have promising potency to rapidly alleviate depressive-like symptoms in preclinical studies (23–26) and emerging clinical evidence supports its good tolerability in humans as well as therapeutic potential to alleviate depressive symptoms (27, 28). LAC is an endogenous short-chain acetyl ester of free carnitine involved in the transport of long chain fatty acids into the mitochondria for degradation by beta oxidation thus, contributing to energy metabolism (29). In addition, LAC can facilitate the removal of oxidative products, provide acetyl groups for protein acetylation, be used as a precursor for acetylcholine, or be incorporated into neurotransmitters such as glutamate, glutamine and GABA (29). However, it is not known whether LAC treatment can counteract brain metabolic alterations specifically observed in the context of stress-induced depression.

In this study, we investigated the ability of LAC supplementation to protect vulnerable mice against stress induced depressive-like behaviors. As indicated above, socially dominant C57BL/6J mice were at higher risk of developing depression-like behavior when exposed to CSDS (8). Here, in order to exclude that the identified vulnerability is a mere reflect of the social stressor used (15), a first aim of this study was to assess the link between social rank and vulnerability to develop depressive-like behaviors using chronic exposure to a non-social (e.g., physical) stressor. To this end, and given that lipid peroxidation has been shown to be increased by restrain stress in the striatum (30), we exposed mice to the well-established 21-day restrain stress protocol (24, 31). Subsequently, we studied the impact of LAC treatment coinciding with the last week of stress exposure on the concentration of up to 20 metabolites in the NAc using *in vivo* ^1^H-MRS at 14 Tesla (20, 32, 33). We then tested mice for depressive-like behaviors, including motivation to explore social conspecifics and coping responses in the FST (34). Besides the evident translational potential of such technique in identifying biomarkers, rodent ^1^H-MRS studies at ultra-high field bridge a potential pathological indicator and its associated molecular signature to physiological mechanisms.

## Methods and Materials

### Animals

Male C57BL/6J mice were used throughout the study (see Supplemental Methods for details).

### Experimental design

One week after arrival, mice were tested for their anxiety and locomotor behaviors in an elevated plus maze (EPM) and open field (OF) (Figure 1A). After four weeks of cohabitation, a social confrontation tube test (SCTT) was used to reveal individual ranks within the home cage tetrad (32). Subsequently, one group of the animals was subjected to a chronic restraint stress (CRS) protocol for 21 days, while the remaining non-stressed animals were submitted to daily handling and body weighting. The impact of chronic stress on behavior was tested in the social behavior test (SB) test (CRS day 20). The experiment performed to investigate the ability of acetyl-L-carnitine (LAC) treatment on behavioral and metabolic outcomes of CRS, included an additional group treated with LAC from CRS day 15. In addition to the SI test, animals were tested in forced swim test (FST) (CRS day 21). Subsequently, ^1^H-MRS was performed at the end of the protocol (day 22).

**Figure 1.**
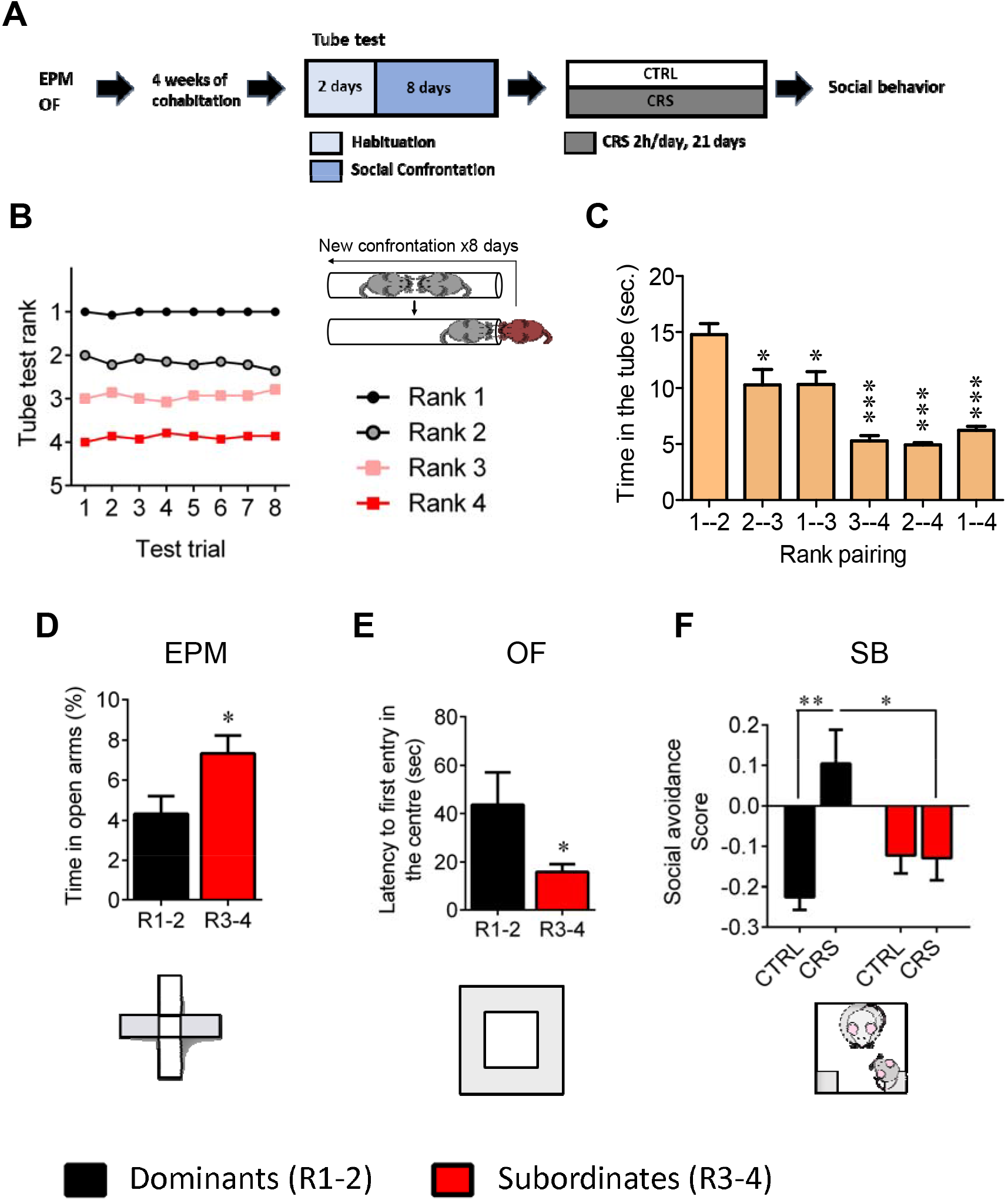
Dominant mice exhibit susceptible behavioral phenotype after 21 days of chronic restraint stress. (A) Experimental design of the restraint stress protocol. (B) Summary of nine cages representing the tube test ranks and winning times as a function of tube test trials over the 6 days of test. (C) Time spent in tube as a function of rank pairing (F_5,35_=18.19, p<.0001, one-way ANOVA; *p<.05, ***p<.001, Bonferroni’s test, n=7 per rank pairing). (D) Trait-anxiety measured based on the time spent in the open arms of an elevated plus maze (p*<.05, unpaired t-test, two-tailed n = 14 per group). (E) Trait-anxiety measured based on the latency to first enter the center of the open-field arena after segregation into dominant vs subordinate mice (*p<.05, unpaired t-test, two-tailed n = 14 per group). (F) Social avoidance scores measured after chronic restraint stress protocol in dominant vs subordinate animals (Interaction: F_1,21_=7.75, P<.05; stress effect: F_1,21_=1.18, P>.05; rank effect: F_1,21_=7.15, P<.05, two-way ANOVA; p*<.05, **p<.01, Bonferroni’s test, n=6-7 per group). Data are displayed as mean ± SEM.

### Behavioral profiling

Anxiety and exploration were tested in the elevated plus maze (EPM) and open field tests (OF), as previously described (35), and social rank established through the social confrontation tube test (SCTT) as detailed in the Supplemental Methods.

### Chronic restraint stress

This protocol involved 21 days of chronic retrain stress (CRS) and was adapted from (24, 31) as detailed in the Supplemental Methods.

### Behavioral testing

Following CRS, animals were tested in the social behavior test (SB) and a social avoidance score was calculated as previously described in (32). They were also tested in the forced swim test (FST). See descriptions of these experimental procedures in the Supplemental Methods.

### Composite behavior

Using MATLAB (Version 9.6, The MathsWorks Inc, Natick, MA), a behavioral composite z-score was calculated by averaging the z-score of the social avoidance score in the SB test and the z-score of the % time spent immobile in the FST. Both z-scores were calculated with the MATLAB function *normalize*, using the option argument zscore, which divides the difference between each sample and the sample average by the sample’s standard deviation.

### Acetyl-L-carnitine treatment

LAC was purchased from (Sigma Aldrich). Mice received LAC in the drinking water at a concentration of 0.3 %. The treatment started on day 15 of the 21 day-CRS protocol and continued till the end of the experiment. Control groups received regular tap water available *ad libitum*.

### ^1^H-magnetic resonance spectroscopy

*In vivo* spectroscopy experiments targeting the Nucleus Accumbens (NAc) were performed in anesthetized mice as previously described (32). Animals were monitored for body temperature (rectal probe and circulating water bath) and respiration (small animal monitor system: SA Instruments Inc., New York, NY, USA) under 1.3-1.5% isoflurane anesthesia mixed with 50% air and 50% O_2_. Physiological parameters were maintained at 36.5 ± 0.4°C and breathing rate ranged between 70 – 100 rpm. Animals were scanned in a horizontal 14.1T/26 cm Varian magnet (Agilent Inc., USA) with a homemade ^1^H surface coil. A set of Fast Spin Echo (FSE) images of the brain was acquired for localizing the Volume of Interest (VOI) of the ^1^H-MRS scan. Acquisition was done using the spin echo full intensity acquired localized (SPECIAL) sequence (36) in the VOI of 1.4 × 4.1 × 1.2 mm^3^ (TE/TR = 2.8/4000ms) including the bilateral NAc after field homogeneity adjustment with FAST(EST)MAP (37). The obtained spectra (20 × 16 averages) were then frequency corrected, summed and quantified using LCModel (38). Full width at half maximum (FWHM) was used as the output of the LCModel. Concentrations were referenced to the water signal and fitting quality assessed using Cramer-Rao lower bounds errors (CRLB) (39). A CRLB value below 20% was used as cutoff for high concentration metabolites, while low concentration metabolites were not considered reliable above a CRLB of 50%.

### Statistics

All values are represented as mean ± SEM. Results from elevated plus maze (EPM) and open field (OF) were analyzed using unpaired Student t-tests. Results from the social confrontation tube test were analyzed with one-way analysis of variance (ANOVA), with social rank as fixed factor, followed by a Bonferroni corrected post hoc test when appropriate. Behaviors in the social behavior (SB) and forced swim (FST) tests were analyzed with a two-way ANOVA, using stress and social rank as fixed factors. In the LAC experiment, behavioral and spectroscopy results were analyzed using one-way ANOVA. Cumulative weight gain was analyzed using a repeated measure two-way ANOVA with time and group as fixed factors. Analyses were followed by Bonferroni post hoc correction when appropriate. Correlations analyses were performed using a Pearson’s correlation coefficient. All statistical tests were performed with GraphPad Prism (GraphPad software, San Diego, CA, USA) using a critical probability of p < 0.05. Statistical analyses performed for each experiment are summarized in each figure legend indicating the statistical test used, sample size (‘n’), as well as degree of freedom, F and P values.

### Factor analysis

Factor analysis was used as previously described (32) using IBM SPSS Statistics version 21 to allow statistical tests using the metabolite’s latent variables as dependent variables in NAc. A linear combination of the dependent variables is generated in order to reduce the noise caused by the high number of variables. Missing values were avoided by using mean value imputation before the computation of correlation matrices, to ensure positive definiteness. A total of three factors was chosen for the NAc after analyzing the scree plots, using principal axis factoring. This resulted in a total of variance explained of 52% without rotation and omitting coefficients below 0.4.

## Results

### Dominant mice are vulnerable to chronic restraint stress

Here, we confirmed and followed up on our initial observations (32) by reporting that dominant and subordinate mice, segregated by using a SCTT (Figure 1B and C; time in the tube 1-2 vs 1-4, p<.001), displayed different profile for anxiety-like behaviors under basal conditions. It was reflected by a higher time spent in the open arms of an EPM (Figure 1D; p<.05) as well as an increased latency to enter the center of an OF (Figure 1E; p<.05) in dominant compared to subordinate mice. Importantly, no difference in locomotor activity was observed, as the distance travelled was similar regardless of the groups (data not shown; n.s.). Strikingly, we revealed that the susceptibility to CSDS observed in dominant mice in our previous study (32) can be generalized to a non-social CRS protocol. Hence, dominant individuals are the ones that were susceptible to develop social avoidance towards an unfamiliar mouse after CRS (Figure 1F) while subordinates seemed not being affected (rank effect, F_1,21_=7.15, p<.05; interaction, F_1,21_=7.75, p<.05; Two-way ANOVA).

### LAC treatment partially abolishes stress-induced behavioral vulnerability in dominant mice

Given the emerging evidence indicating a potential therapeutic efficiency of LAC, the acetylated form of carnitine, in the context of depression (see Introduction), we tested whether LAC treatment could counteract the induction of depressive-like behaviors by CRS. In this experiment, mice were exposed to CRS protocol and received concomitant administration of LAC during the last 7 days of the stress period. Animals exposed to CRS displayed a significant decrease in cumulative body weight gain regardless of the social rank that was not counteracted by LAC supplementation (Figure 2B; in dominant: Stress effect F_2,10_=45.0, p<.0001 and Figure 2C; in subordinates: Stress effect F_2,10_= 20.4, p<.001). As we showed in our previous and current studies that CSDS and CRS induce social avoidance only in dominant mice, we wondered whether LAC supplementation could attenuate the effects of chronic stress on emotional behavior. Although LAC treatment failed to reduce social avoidance after CRS in dominant mice (Figure 2D; F_2,13_=0.55, n.s.), it was sufficient to abolish CRS-induced passive copping strategy in a FST (Figure 2E). As expected, no effects of neither CRS nor LAC treatment were observed in subordinates/resilient mice (Figure 2D). Since LAC has been shown to be a mitochondria-boosting supplement, we tested the possibility that LAC would be more efficient in a high energy-demanding test such as the FST. While CRS induced a significant increase in the immobility time in dominant mice (Figure 2E; F_2,12_=5,31, p<.05), LAC supplementation corrected this behavioral phenotype (p<.05). Again, no effects of neither CRS nor LAC treatment were observed in subordinates/resilient mice (Figure 2E; F_2,15_=1.62, n.s.). These experiments suggest that pharmacological enhancement of mitochondrial function by LAC supplementation normalizes behavioral changes in stressed-dominant mice under unescapable adversity. Finally, we computed an overall behavioral composite to integrate deviation from normality considering the variance in both behavioral tests. ANOVA of these data confirmed these differences (Figure 2F; F_2,13_=10.31, p<.005); with an increase in depressive-like behaviors induced by stress (p<.005) in dominant mice and restored by LAC (p<.05). ANOVA for the composite behavioral z-score for the three groups of subordinate mice indicated no significant differences (F_2,15_=0.16, n.s.).

**Figure 2.**
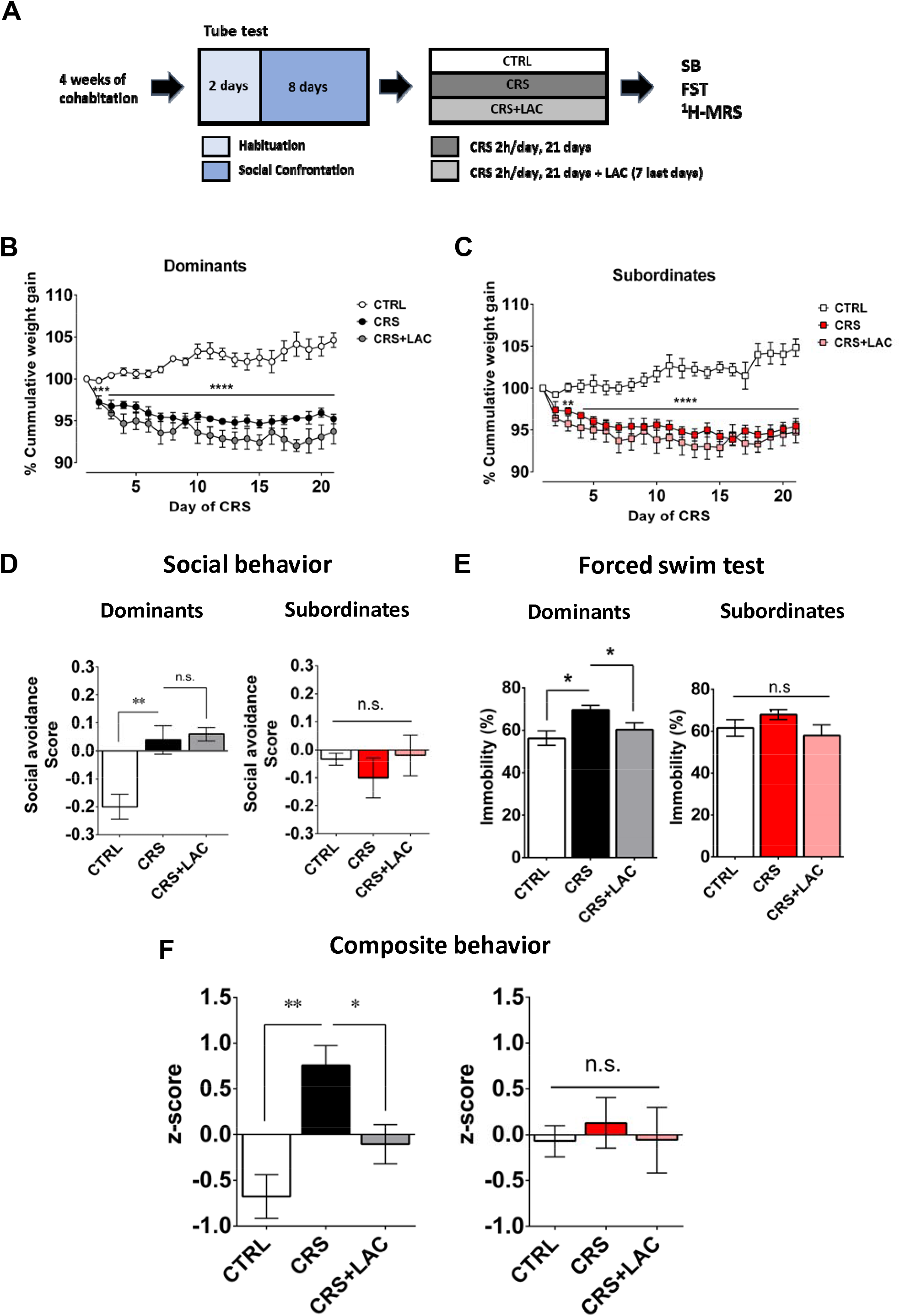
Dominant mice respond to acetyl-L-carnitine treatment after chronic restraint stress. (A) Experimental design of the restraint stress protocol and treatment procedure. (B) Dominant mice show a reduction of cumulative weight gain during the restraint stress protocol (Interaction: F_40,20_=11.5, P<0.0001; stress effect: F_2,10_=45.0, P<.0001, two-way ANOVA; ***p<.001, ****p<.0001, Bonferroni’s test, n=6 per group). (C) Subordinate mice show a reduction of cumulative weight gain during the restraint stress protocol (Interaction: F_40,20_=8.17, P<.0001; stress effect: F_2,10_=20.4, P<.001, two-way ANOVA; ***p<.001, ****p<.0001, Bonferroni’s test, n=6 per group). (D) Social avoidance scores measured after chronic restraint stress protocol in dominant (F_2,12_=12.08, P<.01, one-way ANOVA; **p<.01, Bonferroni’s test, n=6 per group) and subordinate animals (F_2,13_=0.55, P>.05, one-way ANOVA; n=6 per group). (E) Behavioral despair measured with a forced swim test between dominant (F_2,12_=5.31, P<.05, one-way ANOVA; *p<.05, Bonferroni’s test, n=6 per group) and subordinate mice (F_2,15_=1.62, P>.05, one-way ANOVA; n=6 per group). (F) Depressive-like behavior measured as a composite z-score component of social avoidance and immobility time between dominant (F_2,13_=10.31, P<.005, one-way ANOVA; *p<.05**p<.005, Bonferroni’s test, n=6 per group) and subordinate mice (F_2,15_=0.16, P>.05, one-way ANOVA; n=6 per group). Data are displayed as mean ± SEM.

### ^1^H-MRS in NAc reveals stress-responsive metabolites counteracted by LAC treatment

Using *in vivo* ^1^H-MRS, we aimed at revealing NAc neurochemical and metabolite profile in dominants exposed to CRS and supplemented with LAC. The 20 min ^1^H-MRS acquisitions led to a spectral signal-to-noise ratio (SNR) of 17.5±0.3 with a linewidth of 16±1 Hz after shimming with FAST(EST)MAP. The acquired NAc spectra allowed us to quantify up to 20 metabolites with LCModel (Figure 3). First, we applied an unbiased multivariate factor analysis (FA) that revealed 3 main factors that accounted for 31%, 12% and 9% of total variance (Figure 4). Individual metabolites with loadings above 0.4 in factor 1 included taurine (Tau), glutamate (Glu), phosphocreatine (PCr), N-acetylaspartate (NAA), γ-aminobutyric acid (GABA), creatine (Cr), myo-inositol (Ins), aspartate (Asp), phosphocholine (PCho), glucose (Glc), glutathione (GSH) and ascorbate (Asc) (Figure 4A). Metabolites with loading above 0.4 for the two other components were PCho, glycerophosphorylcholine (GPC), and Asc for factor 2, and glutamine (Gln) and lactate (Lac) for factor 3 (Figure 4A).

**Figure 3.**
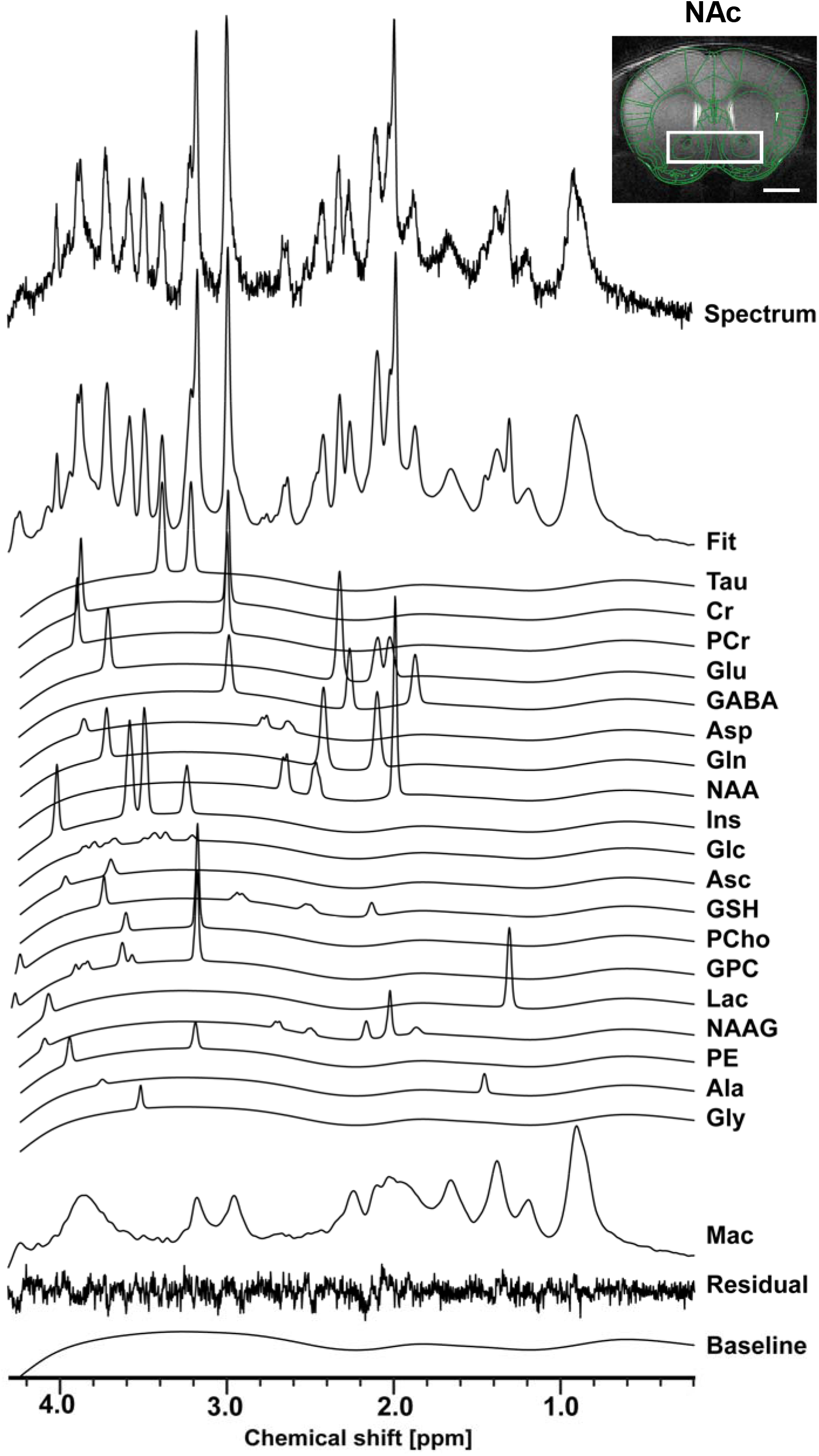
The neurochemical profile of the nucleus accumbens measured with *in vivo* ^1^H-MRS at 14T. (A) Spectrum fitting and neuroanatomical image of the NAc with respective voxel position in mouse brain. Spectrum is decomposed into the total fit, the individual metabolite components of the fit, the residual and the baseline, as a result of LCModel analysis. The fitted neurochemical profile included following metabolites: taurine (Tau), creatine (Cr), phosphocreatine (PCr), glutamate (Glu), γ-aminobutyric acid (GABA), aspartate (Asp), glutamine (Gln), N-acetyl-aspartate (NAA), myo-inositol (Ins), glucose (Glc), ascorbate (Asc), glutathione (GSH), phosphorylcholine (PCho), glycerophorphorylcholine (GPC), lactate (Lac), N-acetylaspartyl-glutamate (NAAG), phosphoethanolamine (PE), alanine (Ala), glycine (Gly), as well as macromolecules (Mac).

**Figure 4.**
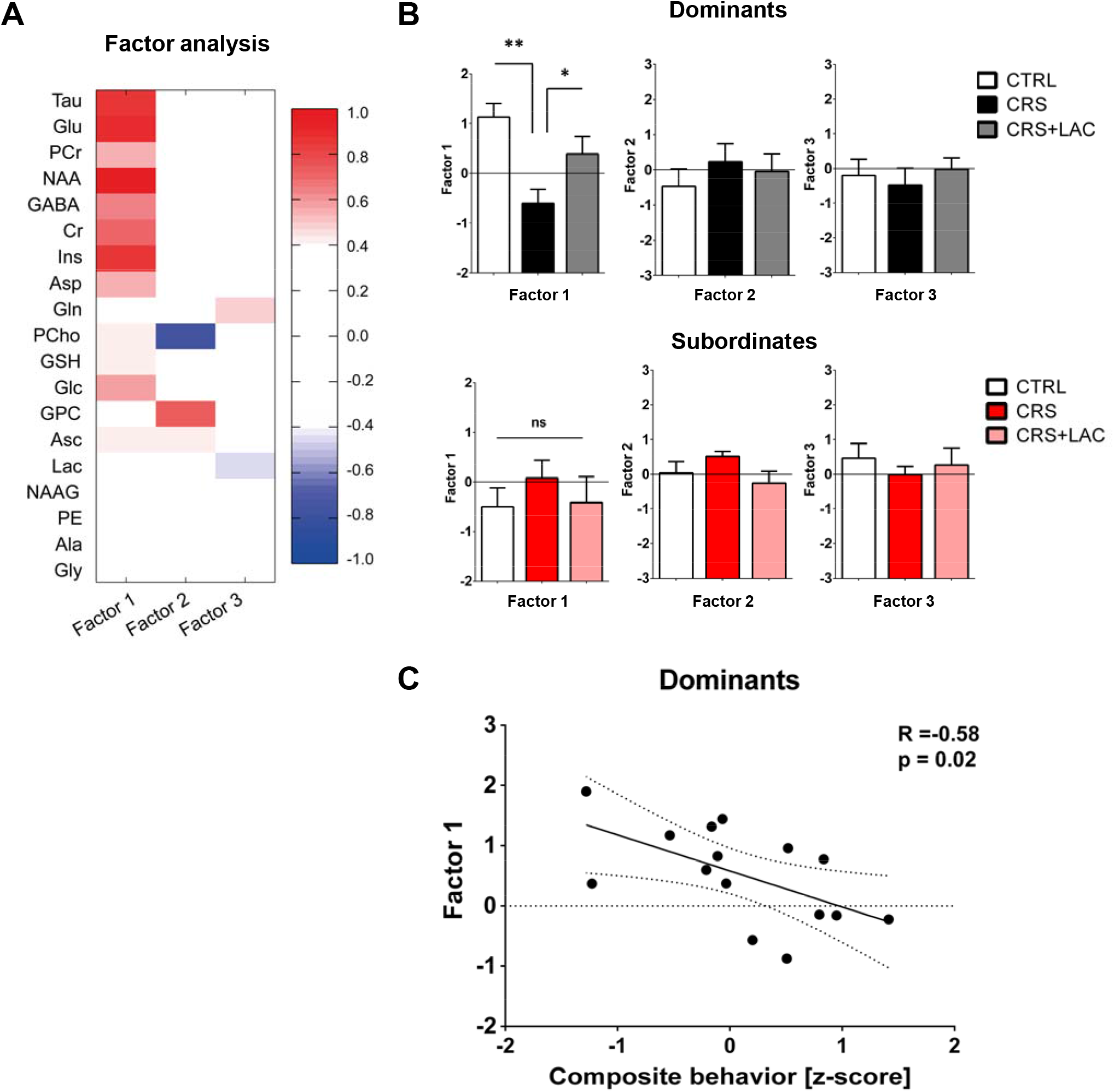
Factor analysis identified one main factor that accounts for the variance in the metabolic profile of nucleus accumbens in dominant mice. (A) Metabolites in the nucleus accumbens that load into Factor 1, Factor 2 and Factor 3. The heat map represents the individual loadings of each metabolite into each factor. (B) Factor 1 represents a linear combination that summarizes neurochemical changes including metabolites with strong (above 0.5: Tau, Cr, PCr, Glc, Glu, GABA, Asp, NAA and Ins.) and moderate (0.4-0.5: GSH and Asc) contribution in dominant mice (F_2,13_=7.04, P<.01, one-way ANOVA; *p<.05, **p<.01, Fisher LSD test n=5-6 per group) and subordinates (F_2,15_=0.54, P>.05, one-way ANOVA; Fisher LSD test n=6 per group). (C) Factor 1 correlates with depressive-like behavior in dominant animals (R=-0.58; p<.05).

Only Factor 1 was able to discriminate for stress and treatment response between dominant/vulnerable and subordinate/resilient mice (Figure 4B; F_2,13_=7.04; p<.01). Hence, CRS led to a reduction in factor 1 metabolites in dominant mice as compared to their controls (p<.005). LAC treatment was sufficient to reverse the effect of CRS on factor 1 (p<.05). Interestingly, we further found a significant negative correlation between the behavioral composite z-score for depressive-like behaviors and factor 1 metabolite levels (Figure 4C; R=−0.58, p<.05).

Then, we selected the metabolites from factor 1 with loadings above 0.4 to perform specific analyses to assess differences between the three groups (i.e., controls, CRS and CRS + LAC) of dominant mice. In metabolites strongly loading in factor 1, we found that the observed stress effect was mainly carried by changes in Tau (p<.05), PCr (p<.05), Glu (p<.), Asp (p<.05) and NAA (p<.05), while the stress-reversing effects of LAC were driven by Tau (p<.05) and PCr (p<.05) (Figure 5). Among the rest of metabolites that either loaded below 0.5 in factor 1 or loaded in the other 2 factors (Supplemental Figure 1S), we found that Gln and PCho were similarly reduced by CRS (p<.05) and reversed by LAC treatment (p<.05). Notably, the ratio of GPC over PCho, that is the level of degradation product of phospholipids over their precursor, respectively, was increased by LAC (p<.05). Overall, metabolite levels in subordinates did not show substantial changes, except for Glc that was increased by CRS (p<.05) and normalized by LAC treatment (p<.05) (Supplemental Figure S2). In further analyses, in an attempt to discover potentially relevant treatment targets that related to animal’s behavior, we performed correlations between each metabolite and behavioral endpoints in data from dominant animals. Two negative correlations were found, one between immobility time in the FST and taurine levels (R=−0.52, p<.05) and a second one between the composite behavioral z-score and GABA (R=−0.55, p<.05) (Supplemental Figure S3).

**Figure 5.**
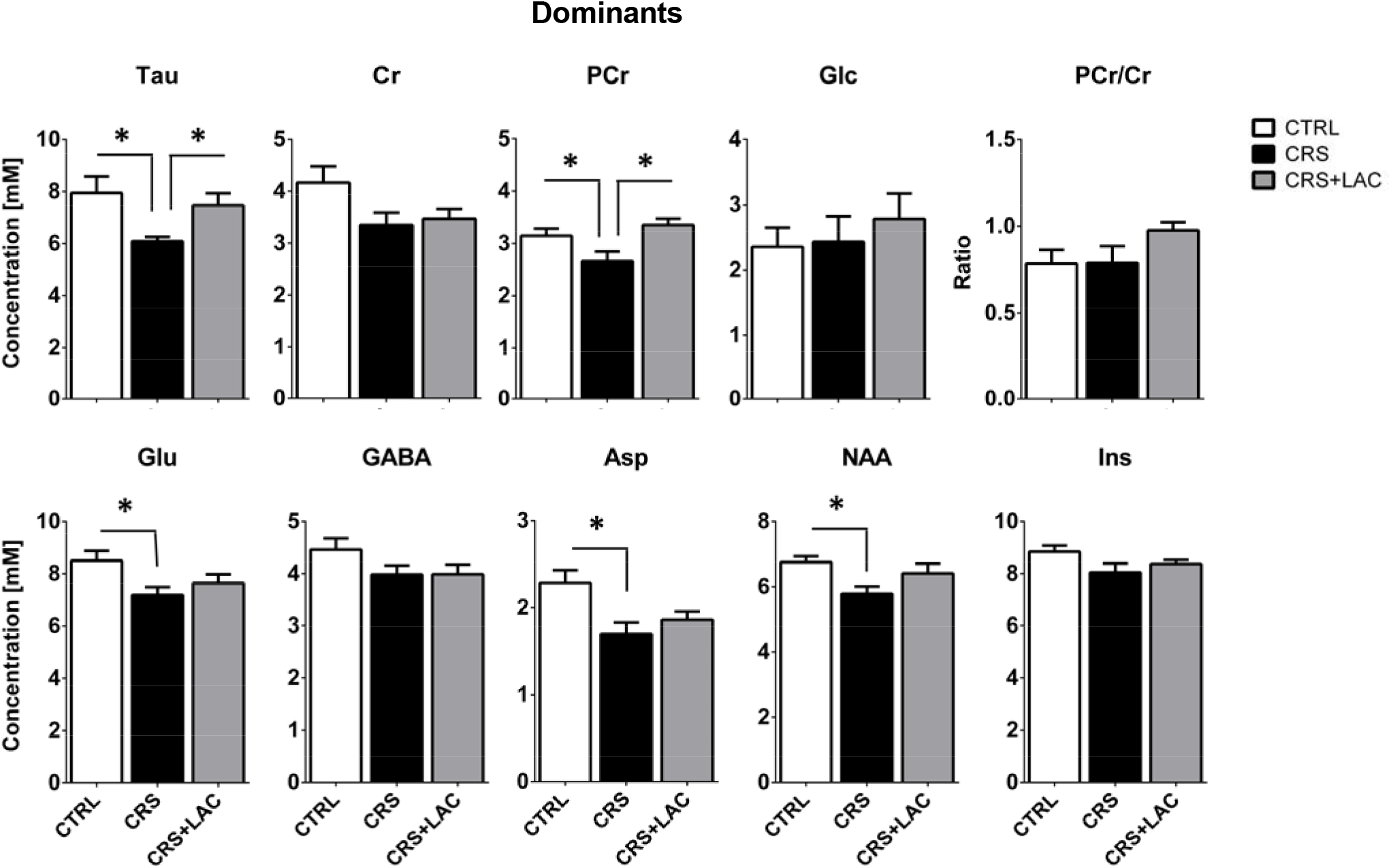
Effect of LAC on the accumbal neurochemical profile of dominant mice after CRS for strong metabolites from factor 1. Metabolites from factor 1 with strong loading (above 0.5) included Tau, Cr, PCr, Glc, Glu, GABA, Asp, NAA and Ins. The ratio of PCr/Cr is shown as well. CRS induces a drop in Tau, Glu, PCr, Asp and NAA but only Tau and PCr are restored after LAC treatment. One-way ANOVA followed by LSD Fisher post-hoc test, *p<.05, n=5-6 per group.

## Discussion

In this study, we first established a link between social dominance in mice and vulnerability to develop depressive-like behaviors following chronic exposure to a non-social stressor. Subsequently, we showed the ability of LAC supplementation to protect dominant/vulnerable mice against a reduction in active coping responses under adversity induced by chronic stress. Furthermore, using *in vivo* ^1^H-MRS at 14 T, we identified a chronic stress-related metabolic signature in the NAc of dominant/vulnerable mice that was partially restored by LAC treatment. Altogether, our results provide an *in vivo* metabolic basis for understanding of antidepressant-like properties of LAC and its protective effects against chronic stress.

When exposed to chronic social defeat (i.e., daily exposure and defeat by an aggressive mouse), socially dominant C57BL/6J mice living in tetrads are at higher risk of developing social avoidance (i.e., a depressive-like behavior) than subordinates (8). Our results reinforce the capacity of social rank in mice to predict vulnerability to stress. in addition, given the non-social character of CRS, our results argue against the view that their vulnerability would be solely based upon loss of social status and resources (15). It is important to note that, as compared to subordinates, dominant mice displayed higher anxiety levels, a well-established risk factor to develop stress-related depressive behaviors (13, 14, 17, 18, 20; for reviews, see Sandi and Richter-Levin, 2009; Russo et al., 2012 and Weger and Sandi, 2018).

Our LAC supplementation regime, administered during the last week of the 3-week CRS protocol, was efficient to protect dominant/vulnerable mice from the development of enhanced passive coping responses (i.e., higher floating levels) in the forced swim test. Importantly, LAC levels are markedly reduced in patients with major depressive disorder, particularly in those with treatment-resistant depression (40). Our results in the forced swim test are in line with previous rodent studies showing the ability of LAC to reduce immobility time in this test (23–25, 41) with several small clinical trials in humans reporting effectiveness of LAC treatment in patients with depressive disorders and related conditions (29, 43, 45–48; for systematic review and meta-analysis see Veronese et al., 2018). Strikingly, LAC supplementation did not change behavioral responses in subordinate/resilient mice that were not affected by CRS, an observation well aligned with findings in the literature indicating lack of LAC effects in non-depressed individuals. Thus, in the genetically-selected Flinders rats, LAC reduces floating behavior in the forced swim test in spontaneously depressed Flinders Sensitive Line (FSL) rats, but it does not change behavior in Flinders Resistant Line (FRL) rats (23, 26). Similarly, no effects of LAC supplementation in this test have been reported for control, unstressed mice (24).

However, the fact that, in our study, the increased levels of social avoidance in chronically stressed dominant mice were not reversed by LAC treatment is at odds with a similar study in which the effects of chronic restrain stress in social behavior were reversed by LAC treatment (24). Although it is difficult to ascertaining the reasons for this discrepancy, they could be due to several methodological differences between the two studies, such as age [mice in our study were slightly older than in Lau et al. (2017)], length of LAC treatment [LAC treatment in our study lasted for 7 days, while only 3 days in Lau et al. (2017)], or mouse substrain [C57BL/6J in our study, while C57BL/6N in Lau et al. (2017)]. In fact, the lack of LAC efficiency in reversing stress-induced deficits in social exploration while its succesful reversal of increased floating in the forced swin test would fit well with a view that the individual’s NAc metabolic *millieu* is critically relevant for the animal’s engagement in energetically-costly behaviors (47). Accordingly, depressive-like behaviors which are more fundamentally dependent on resources mobilization would be more likely to be reversed by LAC’s energy support. Restoring normal metabolic function is, thus, more likely to affect behavior in the context set by the FST – i.e. fighting against an inescapable situation – rather than a social response. This view fits well with the particular focus of this study on energy metabolism in the NAc. Indeed, whereas pharmacological manipulations leading to either impaired or boosted mitochondrial function in the nucleus accumbens were found to boost energetically-costly dominant behaviors during a social competition test between two male rats (47, 48), the same treatments had no effect in animals’ social exploratory behavior (48).

Although different mechanisms can be involved in LAC effects, putative actions through the NAc might be implicated in its capacity to reverse the increase in passive responses induced by chronic stress. For example, active coping in the forced swim test depends upon dopamine actions in the NAc (49, 50) and LAC has been shown to increase DA release *in vivo* (51, 52) and to prevent a chronic stress-induced decrease in DA output in the NAc shell (53). In addition, stress has been shown to induce alterations in the accumbal oxidative stress system (54, 55), and LAC treatment to exert neuroprotective effects by inhibition of glial activation and oxidative stress in the striatum in an animal model of DAergic neuron damage (56). Furthermore, lipid peroxidation has been shown to be increased by immobilization stress in the rat striatum and L-carnitine to reduce associated striatal lipid peroxidation (57). Interestingly, both in rodents (57) and in zebrafish (58), LAC was effective to reverse lipid peroxidation damage in stressed animals while being devoid of effect in controls. These findings resonate with the lack of effect of LAC in subordinate/resilient mice both at the behavioral and metabolic levels (see below).

Our *in vivo* ^1^H-MRS at 14 T identified key metabolites implicated in the response to chronic retrain stress and LAC treatment in the NAc. The PCA factor 1, accounting for 31% of the metabolic variance measured in the NAc, specifically discriminated metabolic responses in the 3 groups of dominant mice: its reduced levels in stressed mice, as compared to controls, were reversed by LAC treatment. Among the 9 metabolites loading highly in this factor, five (i.e., taurine, phosphocreatine, glutamate, aspartate and NAA) were reduced by stress and two of them (i.e., taurine and phosphocreatine) reversed by LAC treatment.

The fact that NAc taurine is identified in our study as critically affected by LAC in the taurine-deficient stressed group is in striking alignment with an earlier human study. Specifically, Sershen et al. (1991) (59) reported that when measuring the impact of a long-term LAC treatment in a human aged population on the levels of amino acids in 5 different brain regions, they only observed a reversal by LAC of ageing-induced reductions in taurine in the striatum, not of any other amino acid in the striatum or of taurine or other amino acids in any other brain region. Taurine is an antioxidant with neuroprotective properties which is located in the mitochondrial matrix and particularly abundant in highly oxidative tissues (60–63). Importantly, in our study, we also found that NAc taurine levels were the only ones from the measured metabolites that negatively correlated with passive coping responses in the forced swim test, reinforcing the link between levels of this amino acid and energetically-costly coping responses to adversity. In a recent 7T ^1^H-MRS in humans, we have recently found a negative correlation between trait anxiety and NAc taurine content (64). Given the high link between trait anxiety and vulnerability to depression (14, 19), our results support the interest in investigating the causal link between NAc taurine and its potential antidepressant actions.

Regarding other metabolites, our findings that LAC increased NAc phosphocreatine levels in dominant/vulnerable stressed mice are consistent with former reports indicating that LAC treatment increases phosphocreatine in the brain (60, 65–68). Accordingly, LAC treatment likely improves the capacity of the brain to produce high-energy phosphates, which may be highly beneficial under conditions of disturbed energy metabolism. Several mechanisms have been implicated in the energy-boosting effects of LAC, many of them relating to an increased oxidative capacity of mitochondria through the direct release of oxidable fuel from LAC itself, or indirectly, in avoiding substrate inhibition of pyruvate dehydrogenase (PDH) by excess of AcCoA (66, 69–71). Our results are in line with the idea of a restored mitochondrial function and support by LAC, visible through the increase in PCr, as well as taurine.

In addition, we observed that LAC restored levels of PCho and the ratio of GPC/PCho, which are disrupted by stress as well in dominant mice. PCho serves as a precursor of phosphatidylcholine (PtdCho), one of the main brain phospholipids, while GPC is its degradation product (72). The GPC/PCho ratio is thus considered to reflect the membrane turnover, typically increased in the case of neurodegeneration or excitotoxicity (73, 74). Increase in GPC can only arise from increased phospholipase activity, which is frequent during excitotoxic events and has been proposed to be a consequence of astrocyte activation (75, 76). LAC and its deacetylated form L-carnitine (LC) are endogenous metabolites involved mainly in the transport and beta-oxidation of lipids. Exchange of LC with LAC and other acylcarnitines trough carnitine-acylcarnitine translocase (CACT) allows a bidirectional flow from cytoplasm into the inner mitochondrial matrix membrane for lipid oxidation. LAC supplementation could thus have a positive effect on phospholipid metabolism, by restoring normal balance between lipid degradation and synthesis.

A role for astrocytes in the LAC mechanism of action is also suggested by the normalization of stress-induced decreases in Gln content observed in the LAC-treated dominant group. Gln is mostly abundant in astrocytes (typically 80% of total concentration) due to their specific expression of glutamine synthetase (GS), which plays a key role in glutamate recycling at the synapse (77, 78). Astrocytic function is fundamental in the resilience to stress and regulation of extrasynaptic glutamate homeostasis (79, 80). For instance, LAC has shown effective control of astroglial cystine-glutamate exchanger (xCT), which is thought to improve mGlu2 function in hippocampus as a response to stress. A similar transcriptional response could be expected for glial GS, given its established responsiveness to glucocorticoids in stress (81, 82), an effect that could underlie the observed Gln changes. Even though the acetyl moiety of LAC has been shown to be utilized for the build-up of metabolic neurotransmitters synthesized from the TCA cycle (83), LAC treatment in our study did not restore the stress-induced decrease of Glu, NAA and Asp observed in the dominant mice. This would suggest that LAC effects on these metabolites may be secondary and part of a slower process. Nevertheless, as NAA, Glu and Asp measured with MRS reflect mainly neuronal metabolism (84, 85), we can hypothesize that astrocytes are the first beneficiary of LAC supplementation. Astrocytes are indeed specifically shaped to uptake blood fatty acids, ketone bodies and acetate, and presumably acetyl-L-carnitine (86–88).

Altogether, our findings highlight an accumbal metabolic signature for vulnerability to stress and response treatment. By implying an accumbal energy- and membrane metabolism process underlying the behavioral outcome, our study identifies molecular candidates responding in opposite direction to chronic stress and LAC treatment, opening possible mechanistic pathways underlying the anti-depressant-like effect of LAC. In particular, we underscore a strong association between NAc taurine and coping behaviors in an energetically-costly adversity task as well as antidepressant LAC actions.

## Acknowledgements and disclosure

This project has been supported by grants from the Swiss National Science Foundation [31003A-152614 and -176206; NCCR Synapsy (51NF40-158776 and -185897)], the European Union’s Seventh Framework Program for research, technological development and demonstration under grant agreement no. 603016 (MATRICS), the EPFL-Jebsen Research Program and intramural funding from the EPFL to CS. The funding sources had no additional role in study design, in the collection, analysis and interpretation of data, in the writing of the report or in the decision to submit the paper for publication. This paper reflects only the authors’ views and the European Union is not liable for any use that may be made of the information contained therein. ^1^H-MRS experiments were also supported financially by the Center for Biomedical Imaging (CIBM) of the University of Lausanne (UNIL), University of Geneva (UNIGE), Geneva University Hospital (HUG), Lausanne University Hospital (CHUV), Swiss Federal Institute of Technology (EPFL) and the Leenaards and Louis-Jeantet Foundations.

The authors declare no financial interests or potential conflicts of interest.

