## Supplemental material for "Metabolic signature in nucleus accumbens for anti-depressant-like effects of acetyl-L-carnitine: An *in vivo* ^1^H-magnetic resonance spectroscopy study at 14 T"

*Supplemental Information*

**Supplemental Methods**

Animals

Six-week-old male C57BL/6J mice were purchased from Charles River Laboratories and, upon arrival, they were housed in groups of four per cage and allowed to acclimate to the animal facility for one week. Mice were weighed at arrival and monitored throughout the experiments. Cages consisted in standard Plexiglass filter-top cages in a temperature (23±1 °C) and humidity (40%) controlled environment with normal 12h day-light cycle. Animals had *ad libitum* access to water and standard rodent chow diet. All experiments were performed with the approval of the Cantonal Veterinary Authorities (Vaud, Switzerland) and carried out in accordance with the European Communities Council Directive of 24 November 1986 (86/609EEC).

Experimental design

One week after arrival, mice were tested for their anxiety and locomotor behaviors in an elevated plus maze (EPM) and open field (OF) (see experimental scheme in Figure 1A). After four weeks of cohabitation, a social confrontation tube test (SCTT) was used to reveal individual ranks within the home cage tetrad (34). Subsequently, one group of the animals was subjected to a chronic restraint stress (CRS) protocol for 21 days, while the remaining non-stressed animals were submitted to daily handling and body weighting. The impact of chronic stress on behavior was tested in the social behavior test (SB) test (CRS day 20). The second experiment, performed to investigate the ability of L-acetyl carnitine (LAC) treatment on behavioral and metabolic outcomes of CRS, included an additional group treated with LAC from CRS day 15. In addition to the SI test, animals were tested in forced swim test (FST) (CRS day 21). Subsequently, ^1^H-MRS was performed at the end of the protocol (day 22).

Elevated Plus maze test

Animals were placed into a maze made from black PVC with a white floor. The apparatus consisted of an elevated central platform (5 x 5 cm^2^) at 65 cm from the ground, from which four opposing arms extended. Two of the arms were open (30 x 5 cm^2^) and lit with 14-15 lx while the two others were closed (30 x 5 x 14 cm^3^) with reduced light intensity 3-4 lx. Animals were introduced in the maze facing the wall at the end of closed arms and left freely moving for 5 min. The mice were video-recorded from above the arena and tracking analyses performed with the Ethovision 11.0 XT software (Noldus, Information Technology) to determine the time spent in open and closed arms.

Open Field test

The OF consisted of a rectangular arena (50 x 50 x 40 cm^3^) illuminated with dimmed light (30 lx). Mice were introduced near the wall of the arena and allowed to explore for 10 min. Analyses were performed using a tracking software (Ethovision 11.0 XT, Noldus, Information Technology) by drawing a virtual zone (15 x 15 cm^2^) in the center of the arena defined as the anxiogenic area. Several parameters were analyzed, including the total distance travelled and the time spent in the different zones.

Social confrontation tube test

The test was performed as previously described (34, 36). Mice were housed together in groups of four during 5 weeks prior to the test to allow enough time for the social hierarchy to be stabilized in the home cage. First, each mouse was independently habituated to cross over a plastic tube (Plexiglas of 3 x 30 cm^2^, diameter x length) on several trials in two consecutive days. Then, evaluation of social rank took place during 8 consecutive days. Specifically, two mice were smoothly guided by the tail to get into each side of the tube for a pairwise confrontation. Once the two mice reached the middle of the tube, the tail was released, and the time spent was recorded until one the mouse (the most subordinate) retracted out from the tube. The four mice from the same cage were opposed using a round-robin design that led to 6 face-to-face trials per day. The tube was cleaned with 70% ethanol and dried after each session. An index of dominance was computed based on the percentage of winning times. This procedure yields ranks which are distributed along a scale from 1 to 4, 1 reflecting the most dominant.

Chronic restraint stress

Animals were introduced head first into a 50ml Falcon tube (11.5 cm in length; diameter of 3 cm) in which the cap was removed. Each restrain tube contained 3 0.4 cm air holes to allow the air to reach the nose of the mouse. Paper was added at the other extremity to adjust the physical constraint to the mouse body size and allowing the tail to reach the open space. The mice were subjected to this restrained environment for two consecutive hours every day for a period of 21 days. Control mice were left undisturbed in their home cage except for handling and body weighting each day for 21 days.

Social behavior test

Each animal was introduced into a 40 x 40 x 30 cm white arena containing an unfamiliar old breeder CD1 male mouse (social target) confined in a cylindrical drum with wire mesh placed near one of the arena walls. The test consists in two phases. First, the experimental mouse was allowed to freely explore for 2.5 min the arena when the social target was absent (the arena contained only the drum). Then, the target mouse was introduced in the drum for a 2.5 min interaction session. A social avoidance score was calculated as previously described in (34). The mice were video-recorded from above the arena and tracking analyses performed with the Ethovision 11.0 XT software (Noldus, Information Technology)

Forced swim test

Each animal was introduced into a cylinder (15 cm diameter, 28 cm in height) filled with 5 L 25°C tap water. The level of water was sufficiently high to avoid any contact of the mouse with the bottom of the enclosure and low enough to avoid any possible escape. Animal’s motion was tracked with a camera positioned on top of the setup and recorded for 6 min. Immobility time was quantified using the Observer XT software (Noldus, Information Technology).

**Supplemental Figures**

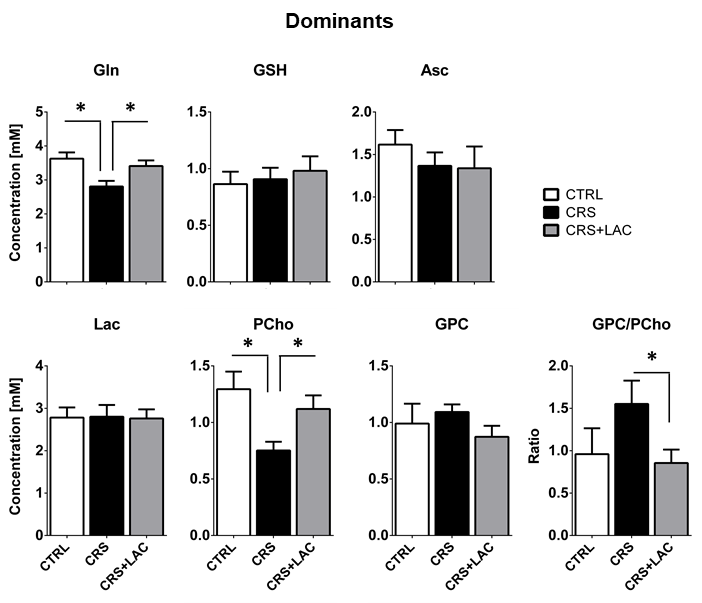

### Figure S1

**Effect of LAC on the accumbal neurochemical profile of dominant mice after CRS for remaining metabolites.** Metabolites with moderate loadings (0.4-0.5) from factor 1 and remaining metabolites from factor 2 and 3 included Gln, GSH, Asc, Lac, PCho and GPC. The ratio of GPC/PCho is also shown. CRS induces a drop in Gln and PCho, which are both restored after LAC treatment. The GPC/PCho ratio is also lowered after LAC administration. One-way ANOVA followed by LSD Fisher post-hoc test, *p<.05, n=5-6 per group.

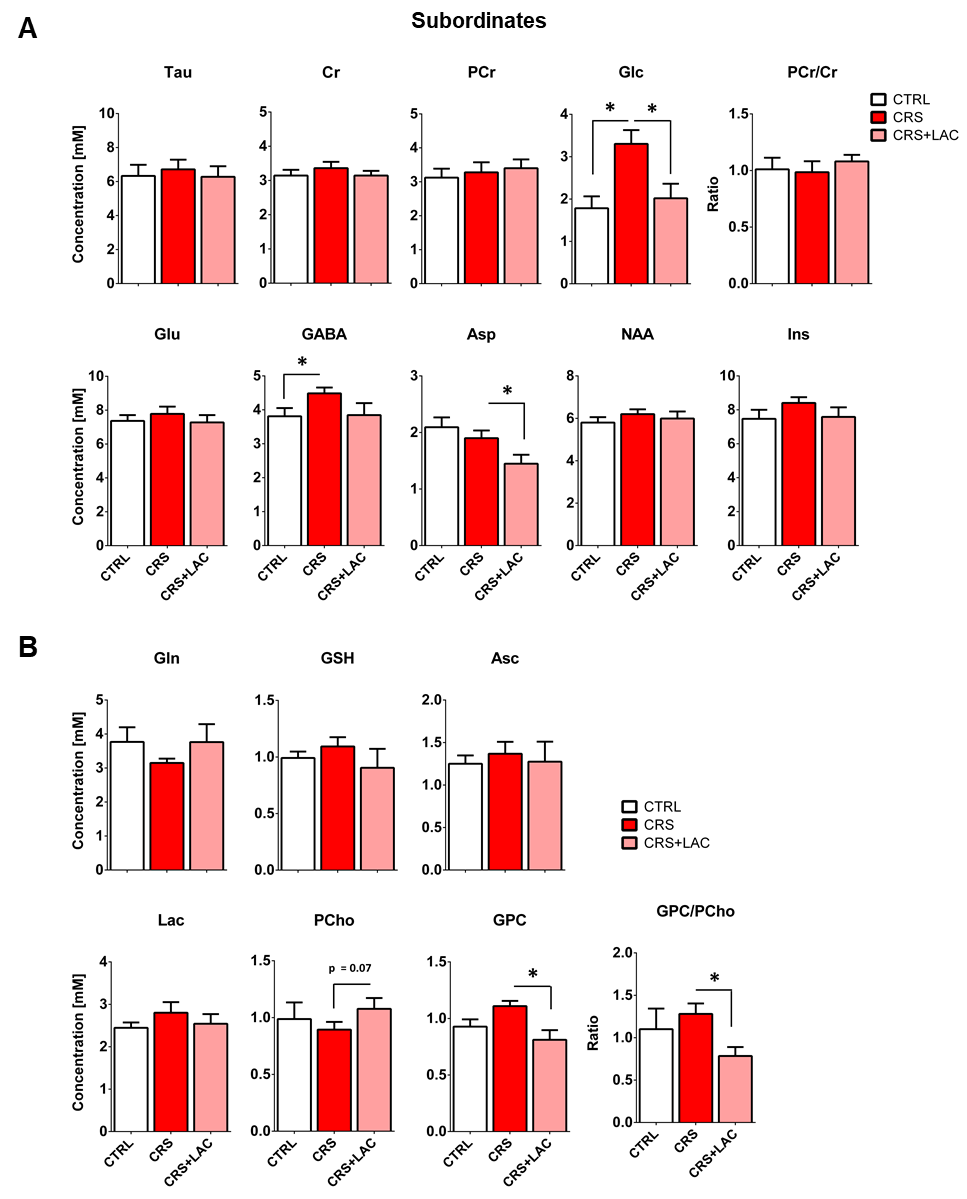

### Figure S2

**Effect of LAC on the accumbal neurochemical profile of subordinate mice after CRS**

(A) Metabolites from factor 1 with strong loading (above 0.5) Tau, Cr, PCr, Glc, Glu, GABA, Asp, NAA and Ins. The ratio of PCr/Cr is shown as well. One-way ANOVA followed by LSD Fisher post-hoc test, n=5-6 per group. (B) Metabolites with moderate loadings (0.4-0.5) from factor 1 and remaining metabolites from factor 2 and 3 included Gln, GSH, Asc, PCho, Lac and GPC. The ratio of GPC/PCho was also reduced upon treatment. One-way ANOVA followed by LSD Fisher post-hoc test, *p<.05, n=5-6 per group.

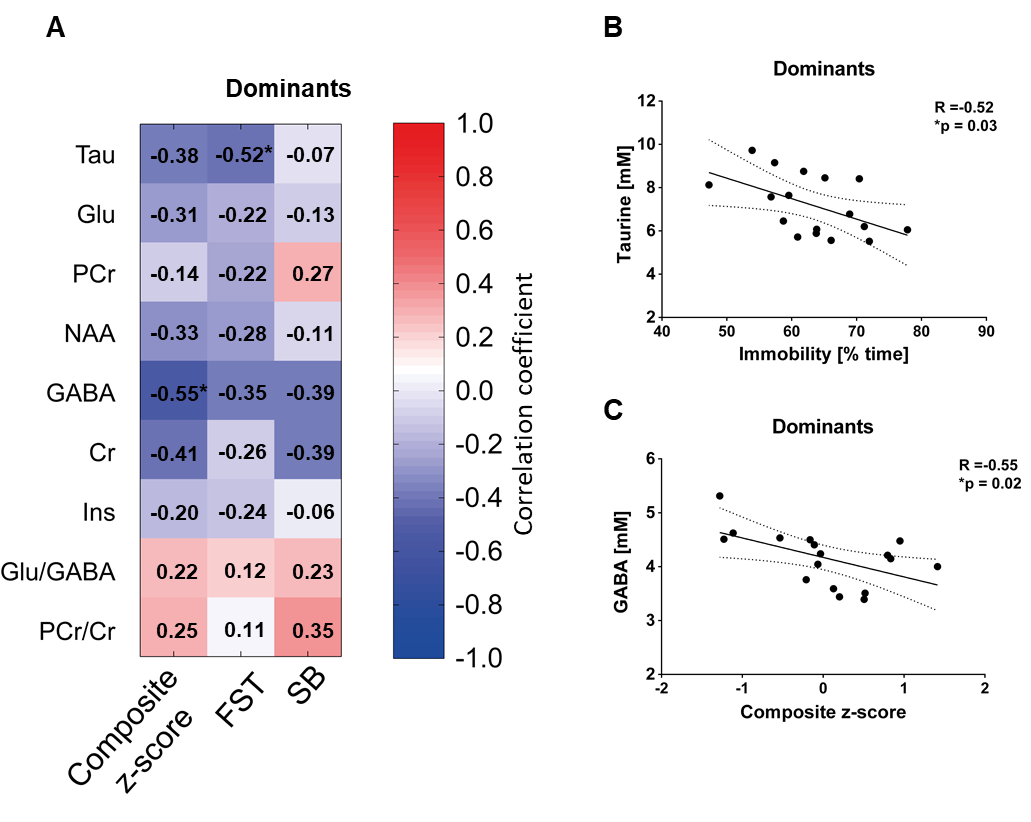

### Figure S3

**Associations between behavior and neurochemistry in the nucleus accumbens.**

(A) Correlation matrix between behavioral components and main metabolic targets of stress in the nucleus accumbens. Behavior included social behavior test (SB), forced swim test (FST) and a composite behavior including both behaviors (Composite z-score). Each cell includes the Pearson’s correlation coefficient with the associated color scaling. (B) Scatter plot of behavioral despair and accumbal taurine. (C) Scatter plot of depressive-like behavior and accumbal GABA. *p<.05, n=16-18 per group.
